# House fly proto-Y chromosomes differentially affect courtship success regardless of social context and female type

**DOI:** 10.64898/2026.09.11.750735

**Authors:** Alireza F. Ghaemmaghami, Rebecca Warneke, Richard P. Meisel

## Abstract

Genetic variation exists for many male reproductive traits despite apparent strong sexual selection in favor of particular males. Understanding how that variation is maintained is an unresolved problem in evolutionary biology. One proposed solution to this problem is genotype-by-environment interactions (GEIs), where no male genotype is universally superior across environments. Another proposed answer is variation in female preference, such that different females prefer different male genotypes. Additionally, male genotypes may differ in their reproductive output as they age. The house fly *Musca domestica* is a well-suited system to evaluate these explanations for genetic variation in male reproductive traits because two male proto-Y chromosome genotypes (Y^M^ and III^M^) differ in mating performance. Specifically, III^M^ males outcompete Y^M^ males in mating assays because III^M^ males have a shorter copulation latency. To determine the relative importance of GEIs, female variation, and male age on the maintenance of genetic variation in male reproductive behaviors, we performed non-competitive (one male to one female) and competitive (one III^M^ male and one Y^M^ male to one female) mating assays while considering multiple ages and different female strains. III^M^ males attempted more matings regardless of competition level, female genotype or age, but mated faster than Y^M^ males only with females of their own strain and only performed more matings in non-competitive trials. However, female strain did not improve the mating success of Y^M^ males. We also did not observe evidence for genotype-by-age interactions on male mating success. Importantly, we did not identify any conditions where Y^M^ males outcompeted III^M^ males, and therefore there was no evidence for interactions or trade-offs that could explain the maintenance of genetic variation for male mating success. To explain the maintenance of III^M^ and Y^M^ males, other factors such as additional social environments or varying temperature should be investigated, as the genotypes are found in natural environments with contrasting temperatures.

## Introduction

Extensive heritable variation exists for male reproductive traits in many animal species, and this variation is often associated with success in courtship and reproduction (Bakker and Pomiankowski 1995). However, if one male genotype is consistently more successful at mating with females or siring offspring, directional sexual selection should increase the frequency of alleles associated with that genotype, eventually eroding the genetic variation underlying male reproductive traits. This raises a fundamental question: how does genetic variation for male reproductive traits persist despite sexual selection?

Answering that question requires considering how male-male competition and female-choice affect the response to sexual selection. Sexual selection can operate through competition among males for access to females and female preferences for particular male traits (Andersson 1994). Female preferences may evolve when females gain either direct benefits (e.g., resources or parental care) or indirect genetic benefits through increased offspring fitness by mating with desirable males (Reynolds and Gross 1990; Kirkpatrick and Ryan 1991). Selection on female preference for indirect benefits requires heritable variation in both male traits and female preferences (Bakker and Pomiankowski 1995). If sexual selection were to reduce genetic variation in male reproductive traits, females would no longer receive indirect benefits from exercising female choice because the progeny of preferred males would no longer attain higher relative fitness (Kirkpatrick and Ryan 1991; Kokko and Heubel 2008). Understanding the maintenance of genetic variation for male sexual traits therefore requires considering variation in female reproductive traits as well.

Many solutions have been proposed that can potentially explain the maintenance of genetic variation in male reproductive traits. One proposed solution to this problem is variation in female mating preferences (Jennions and Petrie 1997). For example, genetic variation in male traits can be maintained if females prefer males with genotypes that differ from their own (negative assortative mating) or simply prefer unrelated males (Hoffman et al. 2007). Secondly, genetic variation can be maintained if genotypes differ in their timing of reproductive investments, such that no genotype maximizes fitness at all ages. For example, the direction of increase in area of coloration, a sexual display indicating male attractiveness in guppies (*Poecilia reticulata*), shows significant family-by-age interactions (Miller and Brooks 2005).

Another proposed solution involves genotype-by-environment interactions (GEIs), whereby the fitness of each male genotype depends on the environment. If the environment varies over space or time, genetic variation can be maintained if no single genotype achieves the highest fitness across all environments (Kokko and Heubel 2008; Ingleby et al. 2010). Genetic variation will therefore be maintained as long as there is gene flow between environmental regimes or across generations experiencing different conditions (Danielson-François et al. 2006; Kokko and Heubel 2008). In the lesser wax moth *Achroia grisella*, for instance, GEIs across different temperatures and densities help maintain variation in male singing traits (Jia et al. 2000). Despite extensive research on the role of GEIs in the maintenance of sexual traits under varying abiotic conditions, little attention has been paid to the role of variation in biotic factors, particularly genotype-by-social environment interactions (Ingleby et al. 2010).

The house fly (*Musca domestica*) is a well-suited model system to investigate the maintenance of genetic variation in sexually selected traits via variation in social environments, GEIs more generally, and other factors. Female house flies seemingly receive no direct benefits from males besides sperm (Carrillo et al. 2012), but see (Arnqvist and Andrés 2006). This limits house fly females to receiving indirect benefits (i.e., offspring with higher fitness because of the paternal genotype), which could contribute to the evolution of female choice (Kirkpatrick and Ryan 1991). Additionally, house flies have a polygenic sex determination system in which the male determining gene (*Mdmd*) is commonly found on two different proto-Y chromosomes, Y^M^ or III^M^ (Sharma et al. 2017; Ronda L. Hamm et al. 2014). Y^M^ and III^M^ are both proto-Y chromosomes that minimally differ from their homologous X and III chromosomes, respectively (Meisel et al. 2017; Son and Meisel 2021). III^M^ males are more likely to mate with females than Y^M^ males when both genotypes are simultaneously presented to a female (Ronda L. Hamm et al. 2009a), likely because III^M^ males tend to have a shorter courtship time (Delclos et al. 2024). Despite the clear mating advantage of III^M^ over Y^M^ males, both genotypes exist in natural populations, even co-existing in some areas (Ronda L. Hamm et al. 2005; R. L. Hamm and Scott 2008). The persistence of both male genotypes in natural populations suggests that other factors may contribute to the maintenance of this polymorphism. Specifically, social factors such as the intensity of male-male competition and variation in female preference could alter the relative mating success of each genotype. Identifying context-dependent effects on male mating performance could therefore explain the maintenance of variation in this sexually selected trait.

Herein, we tested the hypotheses that genetic variation in house fly male mating performance depends on the social environment, female identity, or the age of the mating pair. To those ends, we tested if the mating latency and mating success of the III^M^ and Y^M^ males differ in competitive and non-competitive situations with females from the III^M^ and the Y^M^ strain at different ages. Our experiments allowed us to test whether there are genotype-by-social, genotype-by-age, or male-by-female interactions that could explain the maintenance of both genotypes in natural populations.

## Materials and methods

### House fly strains and rearing protocol

For our experiments, we used a III^M^ house fly strain called CSrab and a Y^M^ strain called IsoCS. Both IsoCS and CSrab share a common genetic background from the CS (Cornell susceptible) strain (Scott et al. 1996), and they only differ in terms of the proto-Y chromosome they carry. The CSrab strain was created by backcrossing the III^M^ chromosome of a spinosad-resistant strain (rspin) from New York (Shono and Scott 2003) onto the CS background (Son et al. 2019). The IsoCS strain was previously created by (Ronda L. Hamm et al. 2009a), from crossing the Y^M^ chromosome of a Maine strain onto the CS background without III^M^. Both the fly larvae and adults were reared and kept at 29℃ with a 12:12 light:dark photoperiod. We used the same larval medium as (Ronda L. Hamm et al. 2009a), and adult virgins were collected within 24 hours of emergence under light CO_2_ anesthesia. We separated the males and females during collection in two mesh cages (30x30 cm) to prevent matings, and we provided flies with *ad libitum* distilled water and a 50:50 powdered milk:sugar mixture.

### Experimental setup

We conducted the experiments with flies aged from 6 to 9 days old to ensure that both males and females were sexually mature (Murvosh et al. 1964). We performed experiments over four consecutive days (Tuesday to Friday), and on a given day all flies were the same age (6, 7, 8, or 9 d old). Within a given week, all flies used in the experiment came from the same cages (described above), without replacement.

We performed two types of assays: non-competitive assays and competitive. Non-competitive assays included one male (either III^M^ or Y^M^) and one female per arena. Competitive assays included two males (one III^M^ and one Y^M^ male) and one female per arena. Females were selected from either the III^M^ or the Y^M^ strain, meaning they were siblings of III^M^ or Y^M^ males and shared the same genetic background but did not themselves carry the male-determining chromosome. The two male genotypes were labeled using yellow and red BioQuip luminous powder by shaking them in an 8 oz paper cup containing the dye, while females were not labeled. The males were labeled in both competitive and non-competitive assays for consistency. The males were then aspirated into 100×25 mm petri dishes (arenas), and females were added after males to prevent matings prior to recordings started. We also added 25×20mm vials to the arenas containing standard *Drosophila* medium media consisting of cornmeal, active dry yeast, sugar, agar, Tegosept, propanoic acid, and water, so the flies could survive for the duration of our recordings.

Within each day, we performed 12 assays, each in its own arena. The twelve arenas were arrayed in four rows of three arenas each and were placed on a white background inside a 29℃ incubator. Our setup was similar to that of (ter Haar et al. 2023), with the addition of two male genotypes (III^M^ and Y^M^) and females from the corresponding III^M^ and Y^M^ strains. For the non-competitive assays on a given day, each arena in the first three rows contained one male and one female, with the male genotypes alternating between the rows. The last row consisted of competitive arenas, with one III^M^ and one Y^M^ male and one female. Across days, we alternated the color assigned to each male genotype and the female strain used in the assays (III^M^ strain or Y^M^ strain females). We performed these assays over the course of six weeks, resulting in 216 non-competitive arenas and 72 competitive arenas. We also ran two weeks of only competitive assays containing 96 arenas with one III^M^ and one Y^M^ male and one female. Within these competitive only batches, the color assigned to each male genotype and the female strain (III^M^ strain or Y^M^ strain females) were alternated between rows.

We used a GoPro HERO 12 camera to record the 12 arenas in each batch. The camera was attached to the ceiling of the incubator. We took images with the time lapse mode every minute for 24 hours. We then manually analyzed the images and scored the behaviors of interest (see below). Any arenas containing dead or escaped flies at any point were dropped from the recordings, resulting in a final total of 162 non-competitive arenas and 141 competitive arenas that were successfully recorded.

### Mating behavior scoring

We scored the following behaviors for each male while analyzing the photos. First, the **number of mating attempts** were scored as male jumps onto females, a behavior termed ‘strike,’ which is the initial step of house fly courtship (Murvosh et al. 1964; Colwell and Shorey 1975). These included any strike that resulted in a male on a female in duration of at least one frame, and it captured both successful and non-successful mating attempts. Second, the **number of successful matings** were scored as copulation attempts lasting at least 30 minutes, which allows for transfer of sperm and most seminal accessory products. Sperm transfer is completed within 10 minutes of copulation (Murvosh et al. 1964), and most seminal accessory products are transferred within 30 minutes (Leopold, Terranova, Thorson, et al. 1971). Third, **copulation duration** was calculated for each successful mating by subtracting the time when the male struck the female until separation. If we observed multiple matings for a male within an arena, we summed the total amount of time each male spent in copula across the matings. We measured copulation duration because housefly male seminal products inhibit females from further matings, and this effect is dose dependent as longer mating durations result in lower frequencies of female remating (Riemann et al. 1967). Fourth, **mating latency** was measured as the amount of time from the start of the recording until the first successful mating.

### Statistical analysis

We analyzed the effects of male genotype, female strain, age of the mating pair, and competition level on each of the four mating behaviors that we measured. For all models, male genotype, female strain, age and competition level were included as fixed effects while week and arena id were included as random effects. For each response variable, we evaluated biologically relevant two-way interactions between male genotype and female strain, competition level, and age. Interaction terms were retained if they significantly improved model fit based on likelihood ratio tests. We fit a zero-inflated negative binomial generalized linear mixed model (GLMM) for mating attempts. Because the data exhibited overdispersion and an excess of zero values (i.e., males that made no mating attempts), a zero-inflated negative binomial distribution provided a better fit than a Poisson model. For the number of successful matings, we fit a Conway-Maxwell Poisson GLMM with a log link because this distribution accommodates both overdispersion and underdispersion and provided a better fit to the data than a standard Poisson model. Copulation duration was analyzed using a GLMM with a Student’s t error distribution after natural log transformation of the response variable because this distribution is robust to deviations from normality and the presence of extreme observations. We analyzed mating latency using a Gamma GLMM with a log link, including only the first mating from trials with successful matings. To test the effect of competition on mating latency, we generated a null distribution of competitive latencies by randomly sampling one III^M^ male and one Y^M^ male from their respective non-competitive latency distributions and calculating the latency to the first mating. The non-competitive distributions included both successful and unsuccessful trials, with a latency of zero indicating that mating did not occur. When both sampled males mated, the shorter latency was used. When only one male mated, its latency was used. If we sampled trials in which neither male mated, we excluded them from the analysis. We then compared these expected distributions with the observed latency to the first successful mating in competitive trials using Monte Carlo permutation tests.

A limitation of the Gamma GLMM analysis of latency is that it requires a successful mating, and arenas without successful matings cannot be included. To overcome this limitation, we also fit a mixed-effects Cox proportional hazards model (treating arenas with zero successful matings as right censored observations) using the coxme package (Therneau 2009). The Cox proportional hazards model included the same fixed and random effects as the Gamma GLMM. All GLMMs were fitted using the glmmTMB package (Brooks et al. 2017) in R version 4.5.2 (R Core Team, 2025). For analysis of each mating behavior, we evaluated model assumptions, including tests for residual dispersion, zero-inflation (where applicable), and residual uniformity using the DHARMa package (Hartig 2016). Additional details are provided in Supplementary File https://doi.org/10.18738/T8/9EVIHL. We additionally evaluated the robustness of significant and marginally significant GLMM coefficients using permutation tests. The resulting permutation-based *p*-values were compared with the corresponding model-based *p*-values, with results summarized in Supplementary Table 1.

## Results

### Mating attempts

We tested how the house fly III^M^ and Y^M^ chromosomes affected the number of mating attempts performed by males in both competitive (two males and one female) and non-competitive (one male and one female) trials. Analyzing the data from both the competitive and non-competitive trials together, Y^M^ males performed approximately 53% fewer mating attempts compared to III^M^ males (*z* = -9.702, *p* = 2x10^-16^) (Figure 1A). There was also a significant effect of female genotype on male mating activity; females of the Y^M^ strain received 26% more mating attempts than females of the III^M^ strain (*z* = 3.015, *p* = 0.0026) (Figure 1A). In some assays, males failed to initiate any mating attempts. There was a marginally significant difference between Y^M^ and III^M^ males in their failure to initiate any mating attempts, with Y^M^ males 2.75 times more likely to not attempt matings (*z* = 1.694, *p* = 0.0902). Both male genotypes had 79% lower odds of not initiating mating attempts in the competitive trials when compared to non-competitive assays (*z* = -2.353, *p* = 0.0186). However, conditioning on males attempting at least one mating, there was no effect of competition level on the number of mating attempts among males that initiated attempts (z = -0.252, p = 0.8007) (Figure 1A). Adding a competition level-by-female strain interaction term resulted in a model with marginally better fit (χ^2^_1_ = 3. 7362, p = 0.0532). In this subsequent model, female strain had no effect on male mating attempts in non-competitive trials (*z* = 0.227, *p* = 0.8201), whereas females of the Y^M^ strain received 36% more mating attempts in competitive trials (interaction: *z* = 1.936, *p* = 0.0529). We obtained similar results when we analyzed competitive and non-competitive trials separately. For example, females of the Y^M^ strain received 38% more mating attempts from males than females of the III^M^ strain in competitive trials (*z* = 3.388, *p* = 0.0007), but not in non-competitive trials (*z* = 0.009, *p* = 0.993).

**Figure 1.**
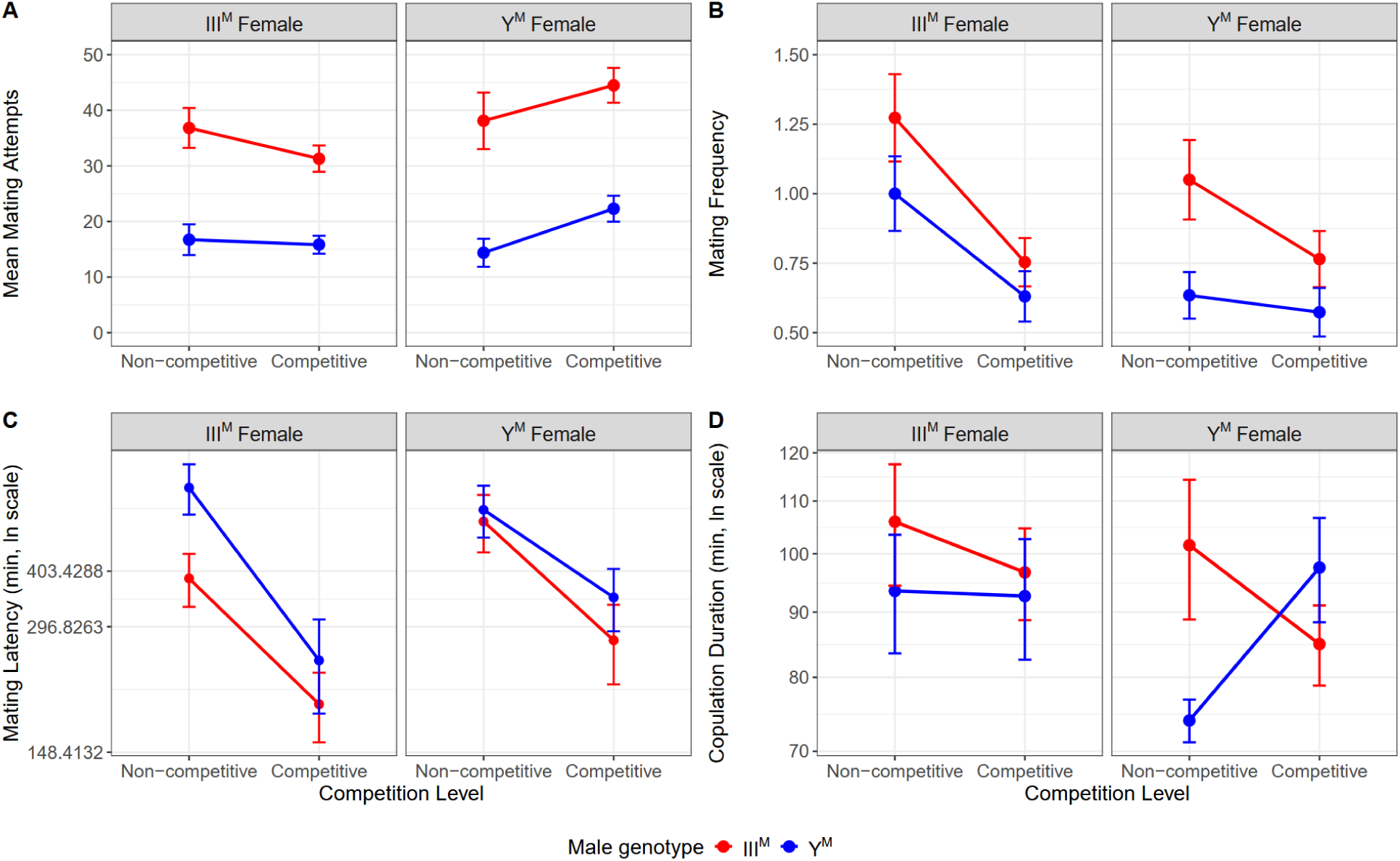
Effects of male genotype, female strain, and competition level on male mating behavior. The Y-axes show the mean **(A)** number of mating attempts per male; **(B)** number of successful matings per male; **(C)** mating latency on a natural log (ln) scale; **(D)** copulation duration per male (± SE) on a natural log scale. Error bars show the standard error of the estimate of the mean. The x-axis indicates competition level, and lines connect treatment means across competition levels for III^M^ (red) and Y^M^ (blue) males. In each panel, matings with females from the III^M^ strain are on the left, and matings with females from the Y^M^ strain are on the right.

Other factors did not affect our results. As examples, adding a male-by-competition level, male-by-age, or male-genotype-by-female strain interaction term to the original model including both competitive and non-competitive trials did not result in models with better fit. We also did not observe a significant effect of age on the number of mating attempts performed by males (*z* = -0.105, *p* = 0.9163) (Figure 2A). We additionally did a separate calculation of *p* values by permutation of our data, and we observed that the same factors significantly affected mating attempts as when we use model-based *p* values reported above (Supplementary Table S1).

**Figure 2.**
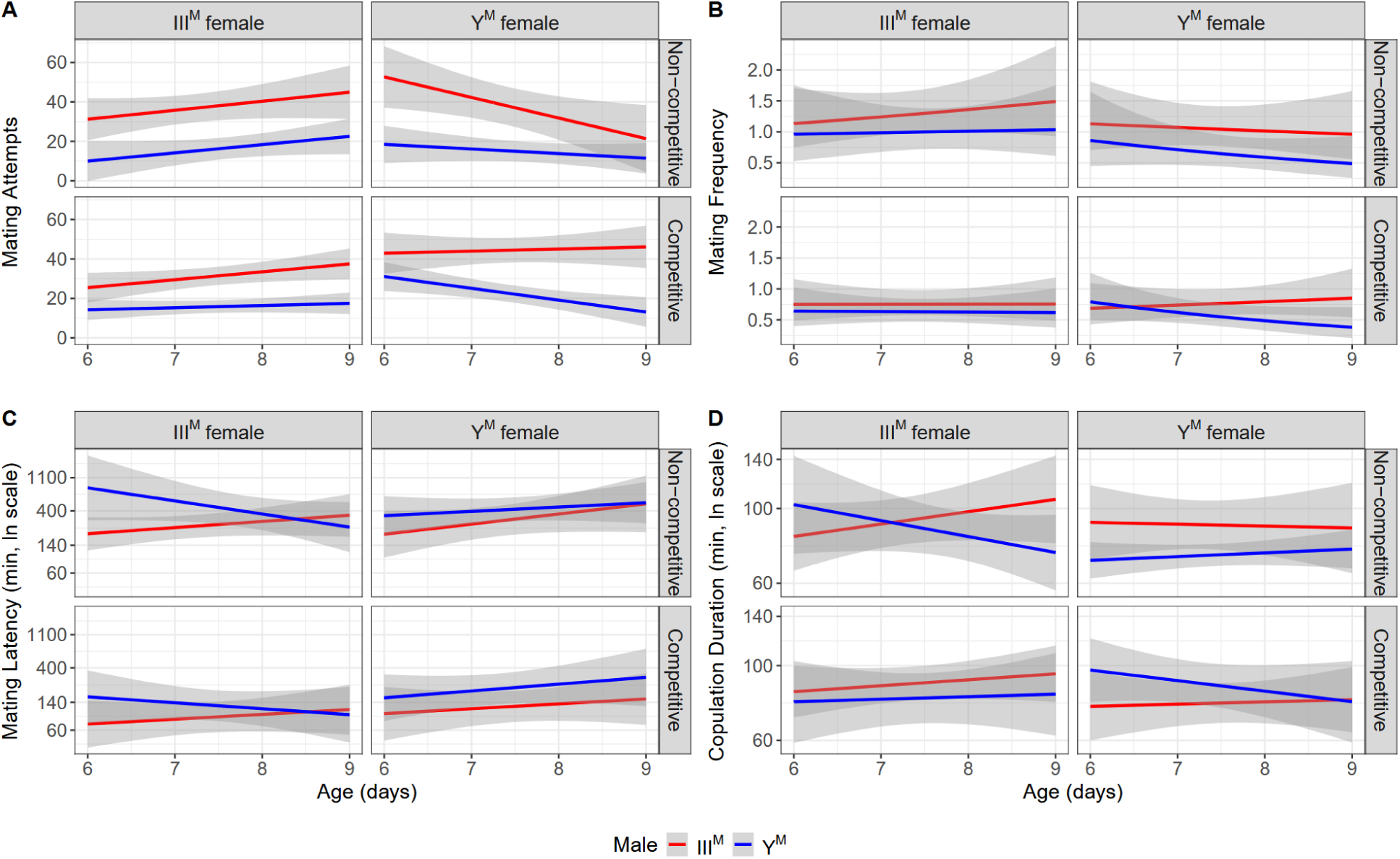
Relationship between male age and mating behavior. **(A)** The linear regression fits for the number of mating attempts are plotted as a function of male age. **(B)** Poisson generalized linear model fits for the number of successful matings are plotted as a function of male age. **(C)** The linear regression fits for mating latency are plotted as a function of male age on a natural logarithmic scale. **(D)** The linear regression fits for copulation duration as a function of male age are plotted on a natural logarithmic scale. For all panels, the x-axis indicates male age in days. Shaded regions represent 95% confidence intervals around the fitted regression models. Panels are separated by female strain (columns) and competition level (rows). Left columns show models fit to matings with females from the III^M^ strain, Right columns show models fit to matings with females from the Y^M^ strain. III^M^ males are shown in red and Y^M^ males in blue.

### Successful matings

We next tested if there was a difference in the number of successful matings between Y^M^ and III^M^ males. Across both competitive and non-competitive trials, Y^M^ males performed 25% fewer successful matings than III^M^ males (*z* = -3.122, *p* = 0.0018) (Figure 1B). Competition level had a significant impact on the number of successful matings, as we observed approximately 35% fewer matings per male in the competitive trials compared to non-competitive trials (*z* = -3.870, *p* = 0.0001). There was a marginally significant effect of female strain on male mating success; males paired with females from the Y^M^ strain achieved approximately 16% fewer successful matings than those paired with females from the III^M^ strain (*z* = -1.856, *p* = 0.0635). We did not detect any age effects on mating success across both competition levels (Figure 2B). We also did not detect a significant male genotype-by-competition level interaction.

To further assess whether the effect of male genotype and female strain differed between competition treatments, we analyzed competitive and non-competitive trials separately. Y^M^ males mated approximately 29% less often than III^M^ males in non-competitive trials (*z* = -2.561, *p* = 0.0104). In competitive trials, Y^M^ males mated approximately 20% less frequently, but this effect was only marginally significant (*z* = -1.754, *p* = 0.0794). Female strain influenced mating success in non-competitive trials, where males paired with females from the Y^M^ strain achieved approximately 30% fewer matings (*z* = -2.703, *p* = 0.0068). There was no effect of female strain in competitive trials (*z* = -0.242, *p* = 0.8086). Inclusion of male genotype-by-female strain or male genotype-by-age interaction terms did not improve the fit of any models. We observed that the same factors significantly affected successful matings when we use model-based *p* values or *p* values from permutations of our data (Supplementary Table S1).

### Mating latency

Among first successful matings, we tested if male genotype and other factors affected mating latency (the time between when a male is first introduced to a female and when they begin to mate). Y^M^ males had a 43% longer mating latency than III^M^ males (*z* = 2.751, *p* = 0.0059). In addition, mating latency was 45% shorter in competitive trials than in non-competitive trials (*z* = −3.739, *p* = 0.0002). Furthermore, mating latency was 27% longer when males were paired with females from the Y^M^ strain, though the effect was only marginally significant (*z* = 1.898, *p* = 0.0577). Adding either a male-genotype-by-competition level, male-genotype-by-female-strain, or a male- genotype-by-age interaction term to the model did not result in a significantly better fit.

Because we observed a marginal female strain effect on latency, we analyzed the data separately for each female type. Y^M^ males exhibited approximately 60% longer mating latencies than III^M^ males when paired with females of the III^M^ strain (*z* = 2.52, *p* = 0.0117). In contrast, there was not a significant difference in mating latency between male genotypes when paired with females of the Y^M^ strain (*z* = 0.91, *p* = 0.3648). The interaction between male genotype and female strain was not significant in the overall model, and therefore these analyses describe male genotype effects within each female strain rather than demonstrating that these effects differed between female strains. We observed that the same factors significantly affected mating latency when we use model-based *p* values or *p* values from permutations of our data (Supplementary Table S1).

We also explored why the mating latency was shorter in competitive trials. Specifically, we asked if shorter latency could be a result of a decreased waiting time until the first mating simply as a result of two males present (i.e., simultaneously observing two independent trials of a Poisson process). Alternatively, shorter latency could be an effect of competition on mating behavior. To test the Poisson model, we generated a null distribution of competitive latencies from the latency times in the non-competitive trials, as described in the Methods. For females from the III^M^ strain, we determined that the Poisson expectation of latency in a competitive trial would be 214 min. In comparison, the observed median latency in competitive trials was 114 min, which is significantly shorter than the expectation (Monte Carlo test, *p* = 0.0006; Figure 3A). We similarly found that the observed median latency for females from the Y^M^ strain in competitive assays (167.5 min) was shorter than the expected (267 min), although this difference was only marginally significant (Monte Carlo test, *p* = 0.0512; Figure 3B). Thus, the latency observed in competitive trials was nearly half the expectation if the two males were independently courting.

**Figure 3.**
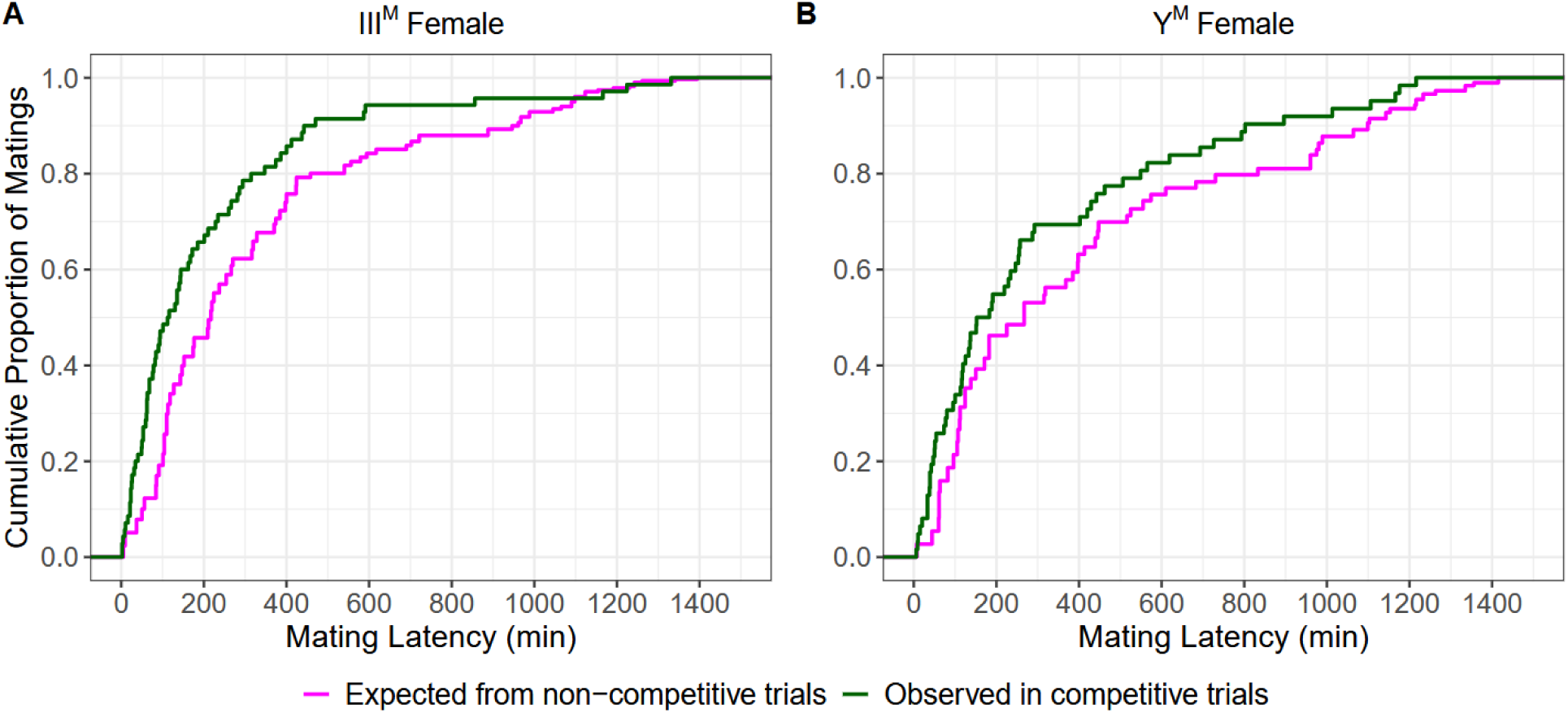
Expected and observed mating latency distributions under competition. **(A)** Mating latency for III^M^ females. **(B)** Mating latency for Y^M^ females. Magenta lines show the expected cumulative proportion of first matings when one III^M^ male and one Y^M^ male are independently sampled from their respective non-competitive mating latency distributions. Dark green lines show the observed cumulative proportion of first matings in competitive trials. Expected distributions include both successful and unsuccessful non-competitive trials. Simulated trials in which neither male mated were excluded.

The linear modeling approach above discards data from unsuccessful matings, which can bias the statistical analysis. To address this limitation, we also analyzed the mating latency data using a Cox proportional hazards (PH) model, which treated unsuccessful matings as right-censored data (Delclos et al. 2024). The hazard ratio (HR) describes the relative instantaneous rate of mating over time, with lower hazard values indicating a lower mating rate at any given time and therefore longer expected time to mating. In a Cox PH model, the Y^M^ males had a lower mating hazard than III^M^ males (HR = 0.61338, *z* = -3.57, *p* = 0.0004), and females of the Y^M^ strain also had a lower mating hazard than those of the III^M^ strain (HR = 0.64875, *z* = -2.87, *p* = 0.0041). Therefore, the Cox PH analysis agrees with the Gamma model for the effects of male genotype and female strain: Y^M^ males had a longer mating latency than III^M^ males, and males paired with females of the Y^M^ strain had a longer mating latency than those paired with females of the III^M^ strain. Unlike the Gamma model, the Cox PH model detected a negative effect of age on mating hazard (HR = 0.82440, *z* = −2.80, *p* = 0.0051), with older flies exhibiting a lower mating rate over time. This difference may reflect the inclusion of unsuccessful trials as right-censored observations in the Cox PH analysis.

### Copulation duration

We found no statistically significant effects of female strain, competition level, age, or any interaction terms on copulation duration (Figure 1C; Figure 2C). However, there was a marginal effect of male genotype on copulation duration, with Y^M^ males copulating for 8% shorter times than III^M^ males (*z* = -1.77, *p* = 0.0774). The permutation analysis provided stronger evidence for a male genotype effect on copulation duration, indicating that III^M^ males mated for longer than Y^M^ males than expected under random reassignment of male genotype (*p* = 0.0009). The shorter copulation duration for Y^M^ males was observed at both competition levels. In non-competitive trials, Y^M^ male copulations were 11% shorter than III^M^ males (*z* = -3.26, *p* = 0.0011). In competitive trials, Y^M^ male copulations were 7% shorter, although this effect was only marginally significant (z = -1.94, *p* = 0.0529). These results suggest that there is a male genotype effect on copulation duration, but competition level does not influence the effect. We similarly observed male genotype significantly affecting copulation duration only in non-competitive trials when using *p* values from permutations of our data (Supplementary Table S1).

## Discussion

We tested how male-male competition, female strain, and age affected the mating performance of two different house fly male genotypes. In nearly every condition, III^M^ males outperformed Y^M^ males. For example, III^M^ males attempted more matings, attained more successful matings, and often had longer copulation durations than Y^M^ males. Rare exceptions to this advantage included trials where matings occurred, wherein III^M^ males did not mate faster than Y^M^ males with females from the Y^M^ strain. Notably, there were no conditions where Y^M^ males outperformed III^M^ males, suggesting that none of the conditions we explored could explain the maintenance of both genotypes.

One reason that the different conditions we tested could not explain the maintenance of both genotypes is that many of the effects were similar for both Y^M^ and III^M^ males. For example, competition influenced several aspects of male mating performance but the effects were in similar directions for both male genotypes. Competition reduced mating latency, with matings occurring sooner when males competed for access to females. Similar reductions in mating latency under competition have been reported in several *Drosophila* species (Bretman et al. 2009; Abraham et al. 2015). While competition did not increase the number of mating attempts performed by males, it substantially increased the probability that males initiated courtship at all. The reduction in mating latency in competitive assays is potentially driven by this greater likelihood of males initiating courtships when a rival is present, thereby increasing the probability of earlier copulation.

Likewise, competition had similar effects on the two male genotypes in terms of mating frequency. Although competition accelerated mating, it reduced the probability that individual males achieved matings as we observed fewer successful matings per male under competition. This reduction in successful matings per male is expected because two males were competing for access to a single female, and female house flies typically mate only once because male seminal products inhibit female remating (Riemann et al. 1967; Leopold, Terranova, and Swilley 1971; Arnqvist and Andrés 2006). Consistent with this constraint, remating occurred in only approximately 20% of our trials. Thus, competition reduced mating opportunities for individual males, but did not affect the two male genotypes differently in mating frequency.

The reduction in mating opportunities under competition could also favor greater investment in each successful mating. In the house fly, male seminal products inhibit female remating, and this effect is dose dependent because longer copulations result in the transfer of larger quantities of seminal products (Riemann et al. 1967). We therefore expected males to prolong copulation under competition, thereby increasing seminal product transfer and reducing the likelihood that females would remate. Although the overall analysis did not support our prediction, we observed a weak trend whereby Y^M^ males appeared to increase copulation duration under competition. This resulted in a smaller difference in copulation times between the two male genotypes in competitive assays when compared to non-competitive trials. However, because we did not detect a significant male-genotype-by-competition level interaction for copulation duration, this pattern should be interpreted cautiously.

Although competition did not significantly alter the relative mating performance of III^M^ and Y^M^ males in our experiments, the smaller difference in mating frequency under competition between the two male genotypes raises the possibility that even stronger competition could reduce the mating advantage of III^M^ males. If the trend we observed were to continue under conditions with even more males, stronger competition could potentially further reduce or even reverse the mating advantage of III^M^ males. Increasingly male-biased sex ratios generally intensify male-male competition (Weir et al. 2011). Accordingly, previous studies in house flies have reported greater overall courtship activity under male-biased sex ratios (Carrillo et al. 2012; ter Haar et al. 2023), although courtship activity per male declines under these conditions ((ter Haar et al. 2023)). However, these studies did not consider the male sex chromosome genotypes, and additional experiments are needed to determine whether III^M^ and Y^M^ males respond differently across a broader range of competitive environments. These assays could potentially reveal scenarios under which the mating advantage of III^M^ males is reduced or reversed.

Female strain represented another factor that could potentially alter the relative performance of III^M^ and Y^M^ males. Similar to competition, female strain influenced male mating behavior, but these effects were generally in the same direction for both male genotypes. Specifically, females of the Y^M^ strain received more mating attempts than females of the III^M^ strain, particularly in competitive trials. Similarly, males paired with Y^M^ females were less likely to mate only in non-competitive trials whereas this difference was not evident in competitive trials. One possible explanation is that females from the Y^M^ strain are more selective than females from the III^M^ strain, mating less readily when presented with a single male but increasing their mating frequency when two males were present. This interpretation is broadly consistent with the increased number of mating attempts directed toward Y^M^ females in competitive trials in our study. Alternatively, Y^M^ females may mate more readily under competition to reduce the costs of persistent courtship and harassment from two males. Previously, it has been shown that in a male-biased sex ratio, female house flies show reduced lifespans as a result of increased male mating activity (Ragland and Sohal 1973). Females could therefore reduce these costs by mating with males. However, because we observed a substantial number of mating attempts directed toward females after they had already mated, the harassment does not appear to be meaningfully reduced by mating.

Female strain also influenced mating latency, although we found no evidence that the identity of the female altered the effect of male genotype on latency. When paired with females of the III^M^ strain, III^M^ males mated faster than Y^M^ males. However, when paired with females of the Y^M^ strain, there was no difference between male genotypes. Because the male-by-female-strain interaction was not significant in the overall model, these results should be interpreted as differences in the male-genotype effect within each female strain rather than evidence that female strain significantly modifies the effect of male genotype. Nevertheless, the absence of a detectable difference in mating latency between male genotypes with Y^M^ females raises the possibility that female responses can influence the differences in male mating performance. These results are similar to a study in *Drosophila pseudoobscura* in which males and females evolved under less competitive (monogamous) and more competitive (polyandrous) sexual selection regimes (Debelle et al. 2016). Males from the polyandrous population outcompeted males from the monogamous population, regardless of female type. However, males from the polyandrous population had shorter mating latencies only with females from polyandrous populations, but not females from monandrous populations.

One possible explanation for the female effect on mating latency in our study is that females may be more willing to mate with males from their own strain. These differences in mating latency could potentially lead to assortative mating. However, the tendency for females to mate more quickly with males from their own strain was insufficient to overcome the overall competitive advantage of III^M^ males, which consistently performed more mating attempts and achieved more successful matings regardless of female type. The basis of the female-strain effect on mating latency is unclear, however, because the strains used in this study share the same genetic background and the females should thus have the same genotype. We therefore suspect that these female strain effects largely reflect differences in the larval environments experienced by the females.

Males and females of the same strain were reared together as larvae and newly emergent adults, which could have influenced female behavior. For example, females may become familiar with the cuticular hydrocarbon (CHC) profile of males from their own strain. There is evidence of CHCs playing important roles in the house fly reproduction (Rogoff et al. 1964; Colwell and Shorey 1977; Adams and Holt 1987). Familiarity of females with the CHC profiles of males from their own strain could potentially contribute to differences in mating performance of the two male genotypes. However, because females of each strain evolved alongside their respective male genotypes during strain construction, the observed female strain effects could also reflect genetic differences that arose during the process of generating the strains. Nevertheless, while female strain clearly influenced several aspects of mating behavior, our results provide only limited evidence that female responses contribute to the mating differences between the two male genotypes.

We considered aging as another potential source of variation in the relative mating performance of the two male genotypes, but found little evidence that age altered their relative performance. The only recorded behavior where we observed an effect of age was mating latency, with older flies exhibiting longer latencies. Because the initiation of copulation in the house fly is under female control (Degrugillier and Leopold 1973), one possibility is that aging effects on female choosiness or mate acceptance contribute to the increased mating latency. Similar age related changes in female mating behavior have been reported in *Drosophila melanogaster* (Travers et al. 2016). We did not, however, observe an increase in mating attempts toward older females, which would be expected if males were to compensate for female age by increasing courtship effort. Alternatively, increased latency in older flies could reflect reproductive senescence in males. Male house fly wings deteriorate more rapidly with age than those of females (Rockstein and Brandt 1963; Ragland and Sohal 1973). Wing buzzing is an important component of male courtship (Meffert and Regan 2002), and wing deterioration caused by aging could reduce males’ ability to perform effective courtship and initiate faster copulations. In addition, the limited age effects observed in our data may simply reflect the relatively narrow age range examined. Specifically, the flies used in our experiments were likely already relatively old. In natural populations, females typically mate at 35–46 degree-days, while males do so at 54–102 degree-days (Butler et al. 2009). The flies in our study ranged from approximately 100 to 150 degree-days old. Experiments involving younger adults would help determine whether age has stronger effects on mating behavior earlier in adult life and whether age interactions with male genotypes occur.

We failed to identify any conditions where Y^M^ males outperformed III^M^ males, but this cannot be attributed to insufficient statistical power. Across multiple measures of mating behavior, we detected consistent effects of male genotype, with III^M^ males outperforming Y^M^ males. We therefore had the power to detect a genotype effect. Our results thus suggest that the lack of evidence for contexts favoring Y^M^ males reflects the consistent direction of the III^M^ male advantage rather than simply an inability to detect a Y^M^ male advantage.

Having found little evidence that competition, female strain, or age provided a context in which Y^M^ males could overcome the mating advantage of III^M^ males, we next considered what might generate this consistent difference in performance. Our results suggest that the III^M^ mating advantage arises from an increase in what we refer to as “courtship effort”. Courtship effort can be considered a form of male-male competition where the male that tries harder wins. We observed that III^M^ males performed more mating attempts than Y^M^ males regardless of mating competition level, female background, or age. Our work also confirmed that III^M^ males have a shorter mating latency than Y^M^ males [(Ronda L. Hamm et al. 2009b; Delclos et al. 2024)]. We hypothesize that the higher courtship effort of III^M^ males explains the larger number of mating attempts, thereby decreasing latency and increasing courtship success in competitive assays. This is consistent with the idea that greater courtship activity can increase mating success (Shamble et al. 2009). However, increased activity can also be associated with fitness costs, including reduced male survival (Kotiaho 2000; Hunt et al. 2004). If III^M^ males suffer greater costs from their higher courtship activity, such trade-offs could reduce the net fitness advantage associated with their superior mating performance. Identifying these trade-offs could explain the maintenance of variation in male mating performance.

In conclusion, our results indicate that the mating advantage of III^M^ males was consistent across the contexts we examined. Although female strain and male competition influenced some elements of mating behavior, these effects were generally not strong enough to reduce the overall mating advantage of III^M^ males. Therefore, variation in female preference and social environment may contribute to the maintenance of both male genotypes, but they are unlikely to do so on their own. Higher competition levels may nevertheless provide contexts in which differences between the two male genotypes are further reduced. Additionally, costs associated with the greater courtship effort of III^M^ males could offset their mating advantage. Identifying the conditions that maintain this polymorphism will therefore require examining these potential trade-offs as well as environmental factors that may differentially affect the two male genotypes, particularly temperature. Natural populations of house flies show a clinal distribution, with Y^M^ males mostly found in the northern latitudes and III^M^ males more abundant in the southern latitudes across different continents (Ronda L. Hamm et al. 2005). In addition, the thermal tolerance and preference of the two types of males is consistent with this geographical distribution (Delclos et al. 2021), and colder developmental temperatures reduce mating latency and increase mating success in both male genotypes (Delclos et al. 2024). Together, these findings suggest that temperature may play a central role in maintaining the house fly sex-determining polymorphism through its effects on mating behavior.

## Acknowledgments

We would like to thank the Meisel lab members, especially the lab manager, Angela Nguyen, for their assistance with house fly husbandry and maintenance in the lab.

## Funding

This material is based upon work supported by the National Science Foundation under Grant 1845686.

## Data availability statement

All data generated in this project and code used to analyze the data are available from the Texas Data Repository at https://doi.org/10.18738/T8/9EVIHL.

## Competing interests

The authors declare that no competing interests exist.

## Notes

https://doi.org/10.18738/T8/9EVIHL

### Competing Interest Statement

The authors have declared no competing interest.

