## Supplemental Table 1 for "House fly proto-Y chromosomes differentially affect courtship success regardless of social context and female type"

***Supplementary Table 1. Comparison of model-based and permutation-based p-values.***

| Response | Effect | Estimate | Model p | Permutation p | Successful permutations | Total permutations |
|---|---|---|---|---|---|---|
| **Mating attempts** | Male genotype | -0.751 | **<0.001** | **0.001** | 998 | 1,000 |
| | Female strain | 0.231 | **0.003** | **0.004** | 1,000 | 1,000 |
| | Competition level (zero-inflation component) | -1.577 | **0.019** | **0.047** | 9,999 | 10,000 |
| | Female strain × competition level | 0.308 | 0.053 | 0.091 | 10,000 | 10,000 |
| | Female strain in competitive trials | 0.322 | **<0.001** | **<0.001** | 1,000 | 1,000 |
| **Successful matings** | Male genotype | -0.287 | **0.002** | **0.002** | 1,000 | 1,000 |
| | Competition level | -0.426 | **<0.001** | **0.002** | 1,000 | 1,000 |
| **Mating latency** | Male genotype | 0.358 | **0.006** | **0.013** | 1,000 | 1,000 |
| | Female strain | 0.239 | 0.058 | 0.075 | 1,000 | 1,000 |
| | Competition level | -0.599 | **<0.001** | **0.003** | 1,000 | 1,000 |
| | Male genotype within IIIM-female trials | 0.473 | **0.012** | **0.025** | 10,000 | 10,000 |
| **Copulation duration** | Male genotype | -0.082 | 0.077 | **<0.001** | 10,000 | 10,000 |
| | Male genotype in non-competitive trials | -0.111 | **0.001** | **0.002** | 1,000 | 1,000 |
| | Male genotype in competitive trials | -0.071 | 0.053 | 0.067 | 9,954 | 10,000 |

*Permutation p-values are two-sided Monte Carlo p-values based on random reassignment of the focal predictor within experimental blocks. Successful permutations indicate the number of permutation fits that produced a positive-definite Hessian out of the total number attempted.*
